# Hypoxia potentiates the inflammatory fibroblast phenotype promoted by pancreatic cancer cell-derived cytokines

**DOI:** 10.1101/2022.07.26.501639

**Authors:** Simon Schwörer, Manon Ros, Kaloyan M. Tsanov, Francesco V. Cimino, Scott W. Lowe, Carlos Carmona-Fontaine, Craig B. Thompson

## Abstract

Cancer-associated fibroblasts (CAFs) are a major cell type in the stroma of solid tumors and can exert both tumor-promoting and tumor-restraining functions. This functional heterogeneity is correlated with the existence of transcriptionally distinct subpopulations of CAFs. CAF heterogeneity is observed in pancreatic ductal adenocarcinoma (PDAC), a tumor characterized by a remarkably dense and hypoxic stroma that features tumor-restraining myofibroblastic CAFs (myCAFs) and tumor-supporting inflammatory CAFs (iCAFs). While CAF heterogeneity can be driven in part by tumor cell-produced cytokines, other determinants shaping CAF identity and function are largely unknown. *In vivo*, we found that iCAFs display a hypoxic gene expression and biochemical profile and are enriched in hypoxic regions of PDAC tumors. Hypoxia leads fibroblasts to acquire an inflammatory gene expression signature and synergizes with cancer cell-derived cytokines to promote an iCAF phenotype in a HIF-1α dependent fashion. Furthermore, we show that HIF-1α stabilization is sufficient to induce an iCAF phenotype in stromal cells introduced into PDAC organoid co-cultures and to promote PDAC tumor growth. These findings indicate hypoxia-induced HIF-1α as a regulator of CAF heterogeneity and promoter of tumor progression in PDAC.

## Introduction

Pancreatic ductal adenocarcinoma (PDAC) is an aggressive tumor and projected to become the second-leading cause of cancer-related mortality by 2030 in the United States (1). A significant barrier to the delivery of effective therapy for PDAC is the desmoplastic stroma that can constitute up to 90% of the tumor volume (1). The prominent desmoplastic response observed in PDAC is characterized by a fibrotic and inflammatory stromal milieu which is produced primarily by cancer-associated fibroblasts (CAFs) and plays a role in both supporting tumor cell growth and promoting therapeutic resistance (2). The basal activity of CAFs to produce extracellular matrix is not sufficient to mediate these effects, as depletion of CAF-derived collagen promotes PDAC growth and reduces survival in mouse models (3). Thus, CAFs can have either tumor-promoting or tumor-suppressing properties within the pancreatic tumor microenvironment (TME).

Transcriptionally and functionally heterogeneous subsets of CAFs have been identified in mouse and human PDAC (4–7). Myofibroblastic CAFs (myCAFs) are marked by expression of alpha smooth muscle actin (αSMA), produce extracellular matrix and are thought to restrain tumor growth (8). Inflammatory CAFs (iCAFs) express only low levels of αSMA, produce a variety of growth factors and inflammatory cytokines such as IL6 and can directly and indirectly promote tumor growth (9). Other, cancer-associated phenotypes of fibroblasts have also been reported, including antigen-presenting CAFs (apCAFs) marked by MHC-II expression (5). Heterogeneity within the CAF population has been suggested to be established in part by growth factor and cytokine gradients within the TME including the local accumulation of tumor-derived TGFβ and IL1/TNFα (10), indicating that spatial differences in the accumulation of different CAF subpopulations exist. However, whether the metabolic conditions present in the pancreatic TME also contribute to regulating CAF heterogeneity is less well explored.

Understanding regulators of CAF heterogeneity has clinical implications: while PDAC patients with high amounts of myCAFs in tumors had improved overall survival, they responded poorly to anti-PD-L1 therapy in retrospective studies (6,8). In contrast, iCAFs are associated with poor response to chemotherapy in patients (11), and iCAF-derived factors including IL6 are directly involved in PDAC progression in mouse models (12–14). Thus, a better understanding of the determinants of CAF heterogeneity may facilitate the development of therapies selectively targeting tumor promoting CAFs.

The TME of PDAC is characterized by nutrient depletion and hypoxia as a result of increased cancer cell demand and impaired vascularization (15,16). Hypoxia results in stabilization of the transcription factor HIF-1α which mediates cellular adaptation to low oxygen tension (17). In cancer cells, this adaptive response promotes epithelial-mesenchymal transition and angiogenesis, and a hypoxia gene expression signature is associated with poor prognosis of PDAC patients (18,19). In the stroma, hypoxia is known to promote lysyl oxidase expression to increase collagen crosslinking and tumor stiffness (20). Hypoxia is associated with an inflammatory fibroblast expression signature in genomic studies of human PDAC and has been shown to promote a secretory phenotype in CAFs while conversely, reducing αSMA expression (21–24). These data suggest that hypoxia could influence the CAF phenotype, but whether hypoxia is involved in the generation of distinct CAF subsets in PDAC is unknown. Here, we report the ability of hypoxia to synergize with cancer cell-derived cytokines to promote the iCAF phenotype and tumor growth in PDAC.

## Results

To investigate factors regulating CAF heterogeneity in PDAC, we analyzed publicly available single cell RNA (scRNA) sequencing data from human PDAC patients (5). Single sample gene set enrichment analysis (GSEA) comparing myCAFs and iCAFs revealed enrichment of an inflammatory response signature in iCAFs and a collagen formation signature in myCAFs (Fig. 1A), as reported (5). Using these data, we found that an oxidative phosphorylation signature was enriched in myCAFs (Fig. 1A), consistent with our previous work showing that mitochondrial oxidative metabolism is required for proline biosynthesis and for collagen production (25). Conversely, a hypoxic gene expression signature was enriched in iCAFs (Fig. 1A). To confirm this finding, we used a murine orthotopic PDAC organoid transplantation model which closely recapitulates key features of human PDAC (26). PDAC organoids derived from the KPC (*Kras*^LSL-G12D/+^;*Trp53*^LSL-R172H/+^;*Pdx1*-Cre) mouse model (27) were injected orthotopically into the pancreas of syngeneic C57BL/6 mice. Once tumors reached ∼500 mm^3^, pimonidazole, a hypoxia indicator (28), was injected intraperitoneally one hour before euthanasia (Fig. 1B). Half of each tumor was digested, and CAFs (gated for CD31^-^CD45^-^EpCAM^-^PDPN^+^ cells) were counterstained for Ly6C as an iCAF surface marker (10) and analyzed for pimonidazole accumulation by flow cytometry (Fig. 1C-E). Consistent with the gene expression data, this analysis revealed that Ly6C^+^ CAFs had accumulated higher amounts of pimonidazole than Ly6C^-^ CAFs (Fig 1D, E). Next, we analyzed the other halves of the PDAC tumors for the presence of pimonidazole, the general CAF marker PDPN and the myCAF marker αSMA by immunofluorescence (Fig. 1F). Strikingly, the vast majority of αSMA^+^ cells was located outside pimonidazole^+^ areas (Fig. 1F). In turn, 80% of the PDPN^+^ areas within pimonidazole^+^ regions stained negative for αSMA (Fig.1F, G).

**Figure 1:**
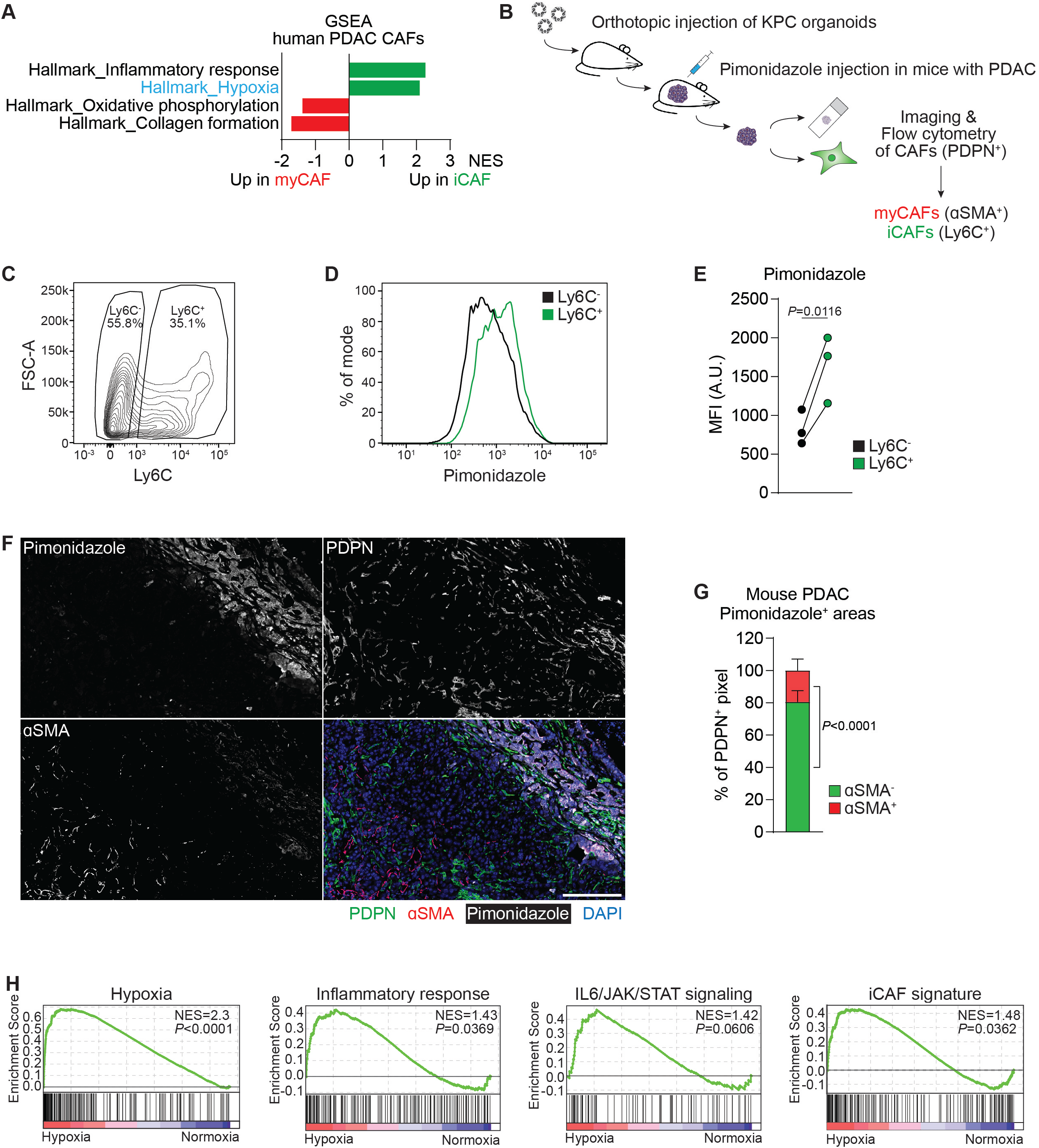
A hypoxic signature is enriched in inflammatory fibroblasts in PDAC. **(A)** Single sample Gene Set Enrichment Analysis (ssGSEA) of selected hallmark signatures in myofibroblastic CAFs (myCAFs) and inflammatory CAFs (iCAFs) based on single-cell RNA-sequencing (scRNA-seq) data from human PDAC. Data from (5). **(B)** Schematic of experimental workflow to analyze Pimonidazole enrichment and localization in mouse PDAC tumors arising from orthotopic transplantation of KPC organoids. **(C-E)** Analysis of Pimonidazole in Ly6C^+^ and Ly6C^-^ cells among live, CD31^-^CD45^-^EpCAM^-^ PDPN^+^ cells in PDAC tumors. **(C)** Gating for Ly6C in PDPN^+^ cells. **(D)** Histogram of fluorescence intensity and **(E)** quantification of Pimonidazole median fluorescence intensity (MFI) comparing Ly6C^+^ and Ly6C^-^ cells. A.U. = arbitrary units. N=3 mice. P-value was calculated by ratio paired t-test. **(F, G)** Immunofluorescence staining of Pimonidazole, PDPN and αSMA in mouse PDAC tumors. **(F)** Representative image. Nuclei are labeled with DAPI. Scale bar = 500 µm. **(G)** Quantification of αSMA^-^ and αSMA^+^ pixel among PDPN^+^ pixel within Pimonidazole-stained regions. N=8 sections from 4 mice. Data represent mean+SD. P-value was calculated by ratio paired t-test. **(H)** GSEA comparing PSCs cultured in normoxia (20% O_2_) or hypoxia (0.5% O_2_) for 48h. iCAF signature derived from (4). Other signatures represent Hallmark signatures from MSigDB. N=2 biological replicates.

The above data indicate a significant positive correlation between hypoxia and the iCAF phenotype in PDAC. To test the hypothesis that hypoxia promotes acquisition of an iCAF state in fibroblasts, we cultured immortalized pancreatic stellate cells (PSCs) for 48 hours in normoxic (20% O_2_) or hypoxic (0.5% O_2_) conditions and interrogated the transcriptome by RNA-sequencing. Hypoxic culture conditions resulted in enrichment of an inflammatory response signature, IL6/JAK/STAT signaling as well as an iCAF signature in PSCs (Fig. 1H).

IL1 and TNFα have been identified as major cytokines secreted by pancreatic cancer cells that are capable of inducing an iCAF phenotype in PDAC (10). In order to assess the role of hypoxia in regulating CAF heterogeneity in relation to known inducers of the iCAF state, we treated PSCs with a combination of IL1 and TNFα (hereafter “cytokines”) to maximize cytokine signalling known to promote an iCAF phenotype. As previously reported (10), cytokine treatment resulted in induction of the iCAF marker *IL6* and repression of the myCAF marker *αSMA* (encoded by *Acta2*, hereafter *αSMA*) (Sup. Fig. 1A). When we cultured cytokine-treated PSCs in hypoxia there was a significant increase in *IL6* expression but no additional changes in *αSMA* mRNA levels (Sup. Fig. 1A). To monitor acquisition of an iCAF state in PSCs by orthogonal methods, we developed a reporter system in which EGFP expression is driven by the murine *IL6* promoter region (Fig. 2A). Responsiveness of the reporter to cytokine treatment was confirmed (Fig. 2A, B). Hypoxia was sufficient to increase the *IL6*-EGFP reporter signal to a similar level as did cytokine treatment, and culture of cytokine-treated cells in hypoxia further increased the reporter signal to more than 15-fold above mock-treated cells cultured in normoxia (Fig. 2A, B). Next, we combined our iCAF reporter with a myCAF reporter in which DsRed expression is driven by the murine *αSMA* promoter region (29). Cytokine treatment reduced *αSMA*-DsRed levels, with hypoxia providing little further reduction of the *αSMA*-DsRed signal at doses of cytokines that maximize *αSMA*-DsRed suppression (Fig. 2C). In contrast, both cytokines and hypoxia increased *IL6*-EGFP levels individually to similar levels and when combined led to marked accumulation of the *IL6*-EGFP signal (Fig. 2C).

**Figure 2:**
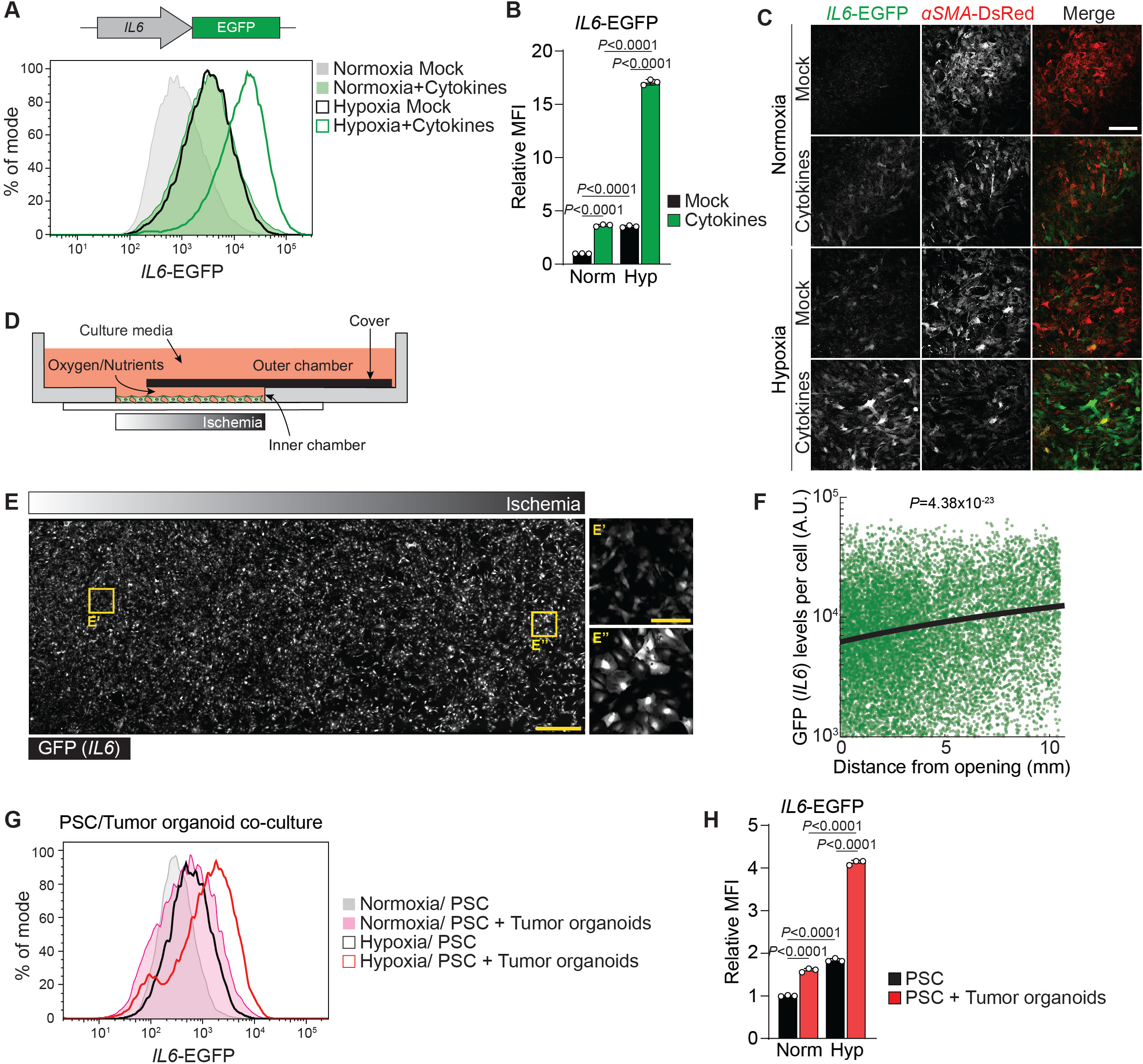
Hypoxia potentiates the cytokine-induced inflammatory fibroblast phenotype. **(A, B)** Fluorescence intensity of *IL6*-EGFP expressing PSCs cultured in normoxia or hypoxia and mock-treated or treated with cytokines (IL1/TNFα) for 48h. **(A)** Histogram of *IL6*-EGFP fluorescence intensity. **(B)** Quantification of the relative MFI of *IL6*-EGFP. N=3 biological replicates. Data represent mean+SD. P-values were calculated by two-way ANOVA. **(C)** Representative images of *IL6*-EGFP and *αSMA*-DsRed expressing PSCs cultured in normoxia or hypoxia and mock-treated or treated with cytokines for 48h. Scale bar = 200 µm. **(D-F)** MEMIC experiment. **(D)** Schematic of the MEMIC, adapted from (31,32). PSCs expressing *IL6*-EGFP were plated in the inner chamber and treated with cytokines the next day. Gradients were allowed to form for 48h. **(E)** Representative image. Cells were fixed and stained for GFP (*IL6*). Nuclei are labeled with DAPI. Scale bar = 500 µM. Oxygen-rich **(E’)** and oxygen-poor **(E’’)** regions are highlighted. Scale bar = 100 µM. **(F)** Quantification of GFP (*IL6*) fluorescence intensity per cell with increasing distance from the oxygen-rich opening. A.U. = arbitrary units. N=15,027 nuclei. Line represents median. P-value was calculated by Pearson’s Linear Correlation Coefficient. **(G, H)** PSC/Tumor organoid co-culture experiment. PSCs expressing *IL6*-EGFP and *αSMA*-DsRed were cultured alone or together with KPC organoids in Matrigel for five days. In the last 48h, part of the cultures was incubated in hypoxia. **(G)** Histogram of *IL6*-EGFP fluorescence intensity in PSCs. **(H)** Quantification of the relative MFI of *IL6*-EGFP in PSCs. N=3 biological replicates. Data represent mean+SD. P-values were calculated by two-way ANOVA.

In tumors, there are gradients of oxygen and nutrient availability (30). To better model these gradients, we cultured PSCs together with cytokines in a metabolic microenvironment chamber (MEMIC) which allows the establishment of oxygen and nutrient gradients within the same culture well (Fig. 2D) (31,32). Using PSCs expressing the hypoxia reporter HRE-dUnaG (33), we confirmed establishment of an oxygen gradient along the MEMIC (Sup. Fig. 1B, C). *αSMA*-DsRed reporter levels gradually declined along the gradient (Sup. Fig. 1B-E). Consistent with the above data, the *IL6*-EGFP reporter signal increased towards ischemic regions (Fig. 2E, F; Sup. Fig. 1D, E), indicating that PSCs acquire iCAF markers in ischemic conditions.

While PSCs cultured on plastic are considered myCAFs, PSCs cultured in Matrigel become quiescent and can acquire an iCAF state when co-cultured with PDAC organoids in Matrigel (4), a process dependent on organoid-derived cytokines (10). Consistent with this, we observed higher levels of *IL6*-EGFP but lower levels of *αSMA*-DsRed in PSCs co-cultured with KPC organoids in Matrigel for five days (Sup. Fig. 2A-D). Next, co-cultures of PSCs and KPC organoids were placed in hypoxia for the last 48h of the culture period. Hypoxia was sufficient to elevate expression of *IL6*-EGFP in PSCs to similar levels as did organoid co-culture, and exposure of co-cultures to hypoxia further elevated *IL6*-EGFP reporter levels in PSCs (Fig. 2G, H).

The above data indicate that hypoxia potentiates the ability of cancer cell-secreted cytokines to promote acquisition of an iCAF phenotype in PSCs. To define the underlying mechanism, we analyzed transcription factor activity in CAFs in human PDAC scRNA-sequencing data by Virtual Inference of Protein Activity by Enriched Regulon (VIPER) analysis (5). As expected, high SMAD2 activity was found in myCAFs, while STAT3 activity was enriched in iCAFs (Fig. 3A). In addition, HIF-1α activity was enriched in iCAFs (Fig. 3A). *Hif1a* is not transcribed basally in resting fibroblasts and its transcription is induced by growth factor and/or cytokine stimulation (34). Even when transcription is induced, fibroblasts like other cells do not accumulate HIF-1α protein due to the oxygen-dependent degradation by VHL (35). Like fibroblasts, PSCs accumulated little HIF-1α under hypoxia, however, when stimulated by cytokines under hypoxic conditions HIF-1α was upregulated synergistically, and we observed increased expression of the HIF-1α target LDHA compared to hypoxia alone (Fig. 3B). Higher levels of HIF-1α were also found in PSCs co-treated with cytokines and cobalt chloride (CoCl_2_), a known inducer of HIF-1α stabilization and signaling (35), compared to CoCl_2_ treatment alone (Sup. Fig. 3A). While CoCl_2_ treatment alone could also increase levels of the *IL6*-EGFP reporter, combined treatment with CoCl_2_ and cytokines elevated the *IL6*-EGFP signal even more (Sup. Fig. 3A-C).

**Figure 3:**
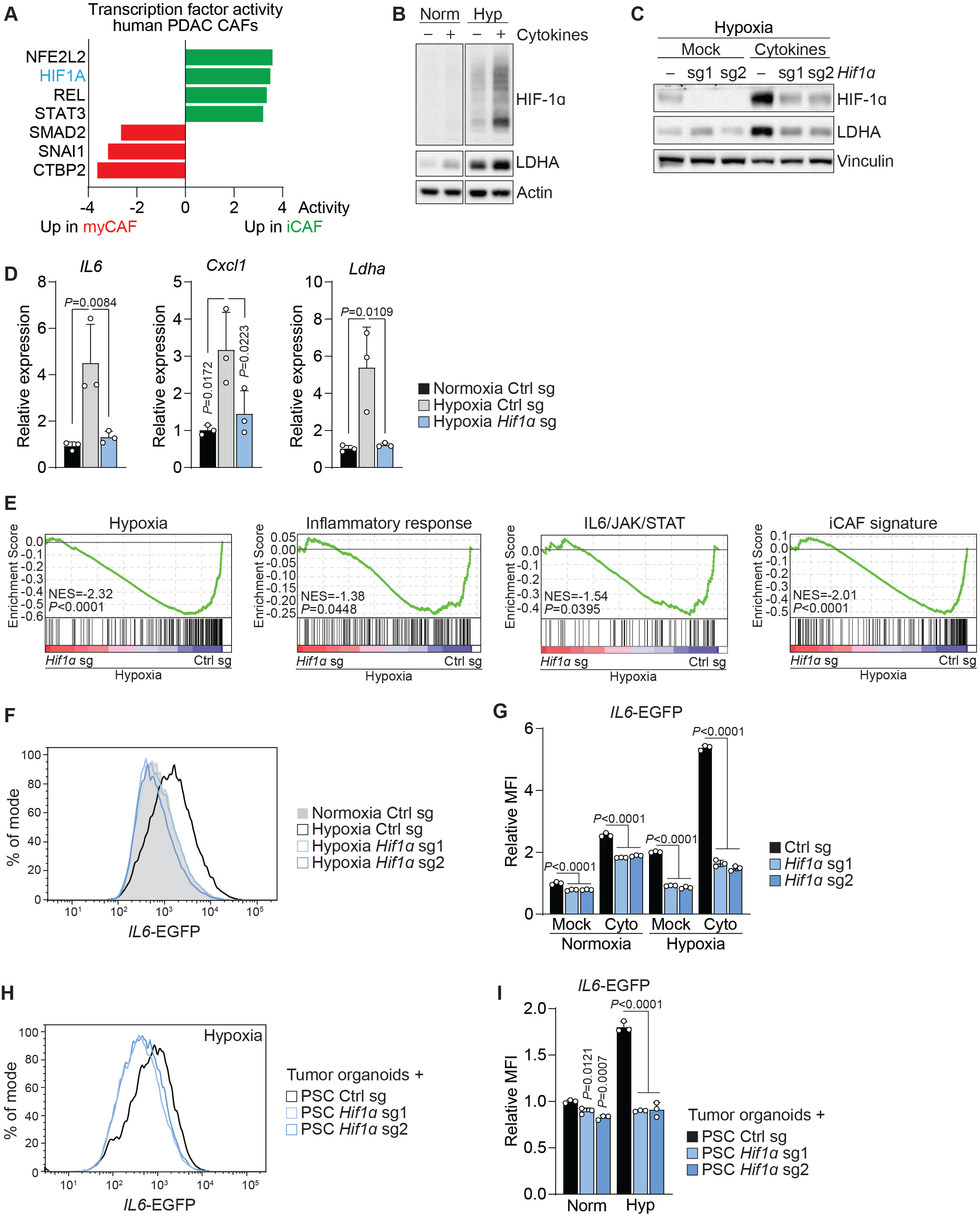
HIF-1α mediates the hypoxia-induced inflammatory phenotype in fibroblasts **(A)** Activity of a selected set of transcription factors in myCAFs and iCAFs based on scRNA-seq data from human PDAC. Data from (5). **(B, C)** Western blots of **(B)** PSCs cultured in Normoxia or Hypoxia and **(C)** PSCs expressing control or *Hif1a* sgRNA and cultured in Hypoxia. Cells were mock-treated or treated with cytokines for 48h. Representative experiments are shown. Separate panels in (B) are from the same membrane with irrelevant lanes cut out. **(D)** qPCR for the indicated transcripts in PSCs expressing control or *Hif1a* sgRNA and cultured in Normoxia or Hypoxia. N=3 biological replicates. Data represent mean+SD. P-values were calculated by one-way ANOVA. **(E)** GSEA comparing PSCs expressing control or *Hif1a* sgRNA and cultured in Hypoxia for 48h. iCAF signature derived from (4). Other signatures represent Hallmark signatures from MSigDB. N=2 biological replicates. **(F, G)** Fluorescence intensity of PSCs expressing *IL6*-EGFP and control or *Hif1a* sgRNA cultured in Normoxia or Hypoxia and mock-treated or treated with cytokines for 48h. **(F)** Histogram of *IL6*-EGFP fluorescence intensity in mock-treated cells. **(G)** Quantification of the relative MFI of *IL6*-EGFP. N=3 biological replicates. Data represent mean+SD. P-values were calculated by two-way ANOVA. **(H, I)** Fluorescence intensity of PSCs expressing *IL6*-EGFP and control or *Hif1a* sgRNA co-cultured with KPC organoids for five days. In the last 48h, part of the cultures were incubated in Hypoxia. **(H)** Histogram of *IL6*-EGFP fluorescence intensity in PSCs cultured with organoids in Hypoxia. **(I)** Quantification of relative MFI of *IL6*-EGFP in PSCs. N=3 biological replicates. Data represent mean+SD. P-values were calculated by two-way ANOVA.

To investigate the role of HIF-1α in regulating the iCAF state in hypoxia, we expressed *Hif1a* sgRNAs which reduced HIF-1α protein levels in mock-treated at well as cytokine-treated cells in hypoxia (Fig. 3C). Induction of *IL6, Cxcl1* and *Ldha* mRNA in PSCs cultured in hypoxia was dependent on *Hif1a* (Fig. 3D). On a global gene expression level, inflammatory response, IL6/JAK/STAT signaling and iCAF signatures were depleted in hypoxic PSCs expressing *Hif1a* sgRNA (Fig. 3E). In addition, the hypoxia-induced increase in *IL6*-EGFP fluorescence required *Hif1a* (Fig. 3F, G). Given the upregulation of HIF-1α in hypoxic cells by cytokine treatment, we also analyzed *Hif1a* sgRNA expressing PSCs in the presence of cytokines. *Hif1a* sgRNA prevented the synergistic accumulation of *IL6*-EGFP in cytokine-treated PSCs cultured in hypoxia (Fig. 3G). Similar results were obtained in *Hif1a* sgRNA expressing PSCs treated with CoCl_2_ (Sup. Fig. 3D). Moreover, the hypoxia-induced upregulation of *IL6*-EGFP reporter levels in PSCs co-cultured with KPC organoids without addition of exogenous cytokines beyond those produced by organoid cultures was also was dependent on *Hif1a* (Fig. 3H, I).

Next, we investigated whether HIF-1α stabilization can be sufficient to shift fibroblasts towards an iCAF state. To induce HIF-1α accumulation under normoxic conditions, we deleted *Vhl*, which targets hydroxylated HIF-1α for proteasomal degradation (35) (Fig. 4A). *Vhl* deleted PSCs displayed higher expression of *IL6, Cxcl1* and *Ldha* mRNA (Fig. 4B). *Vhl* deletion alone increased *IL6*-EGFP levels more than cytokine treatment, and when combined, *Vhl* deletion and cytokines elevated *IL6*-EGFP reporter signals ten-fold (Fig. 4A, C, D). *Vhl* deletion also promoted *IL6*-EGFP signal in PSCs co-cultured with KPC organoids without addition of exogenous cytokines (Fig. 4E, F). Given that iCAFs can promote tumor growth (10), we sought to test whether *Vhl* deletion in PSCs would increase their ability to promote tumor growth *in vivo*. Co-injection of PSCs together with KPC pancreatic cancer cells promoted tumor growth compared to KPC cells alone in a subcutaneous allograft model (Fig. 4G), as reported before (36,37). Notably, co-injection of *Vhl*-deleted PSCs increased tumor growth significantly more than control PSCs (Fig. 4G). Taken together, our data indicate hypoxia-induced HIF-1α as a novel regulator of CAF heterogeneity and tumor growth in PDAC.

**Figure 4:**
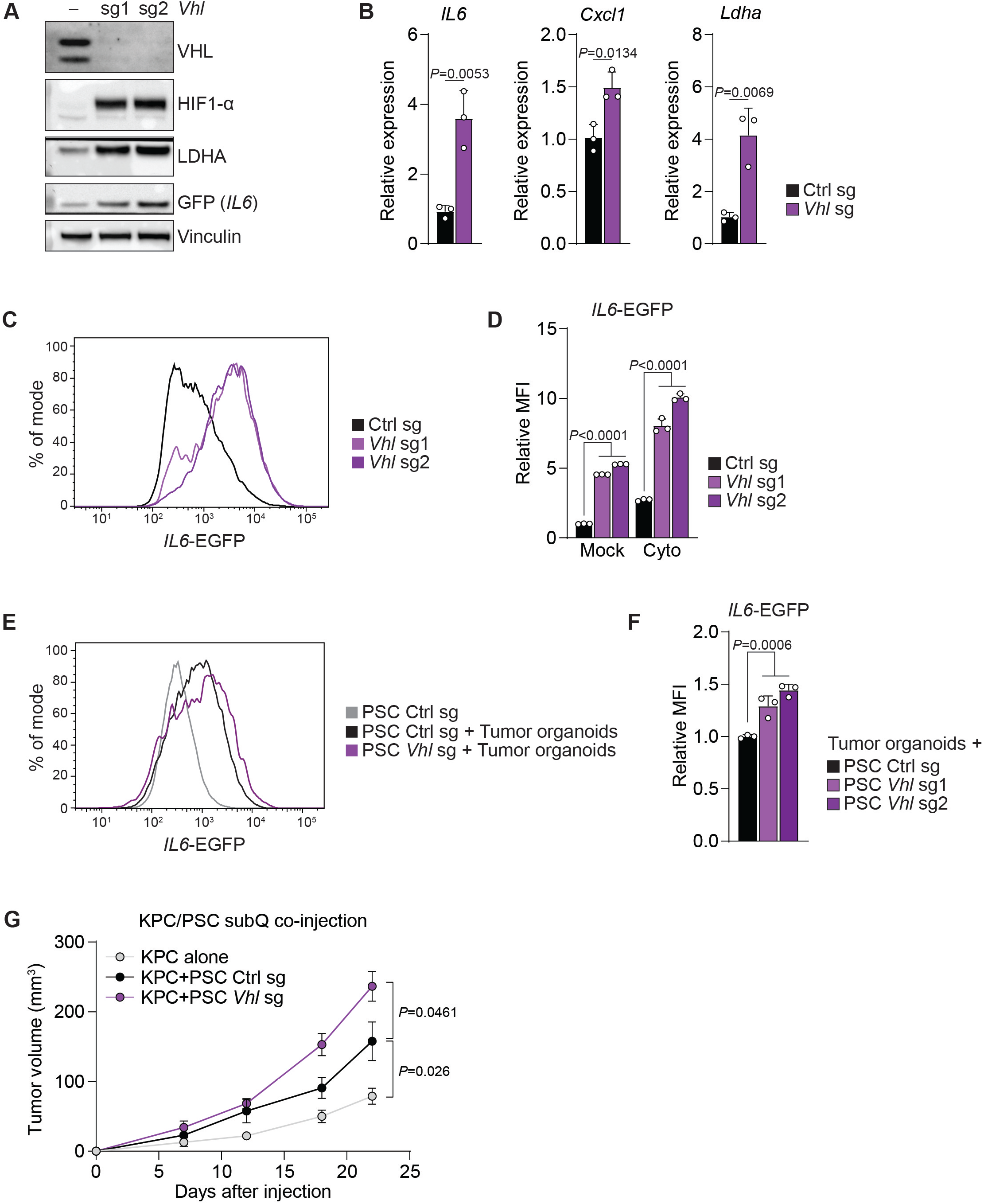
HIF-1α stabilization in fibroblasts can be sufficient to promote an inflammatory phenotype and tumor growth. **(A)** Western blot of PSCs expressing control or *Vhl* sgRNAs and cultured in normoxia. A representative experiment is shown. **(B)** qPCR for the indicated transcripts in PSCs expressing control or *Vhl* sgRNA cultured in normoxia. N=3 biological replicates. Data represent mean+SD. P-values were calculated by Student’s t-test. **(C, D)** Fluorescence intensity of PSCs expressing *IL6*-EGFP and control or *Vhl* sgRNA cultured in normoxia and mock-treated or treated with cytokines for 48h. **(C)** Histogram of *IL6*-EGFP fluorescence intensity in mock-treated cells. **(D)** Quantification of the relative MFI of *IL6*-EGFP. N=3 biological replicates. Data represent mean+SD. P-values were calculated by two-way ANOVA. **(E, F)** Fluorescence intensity of PSCs expressing *IL6*-EGFP and control or *Vhl* sgRNA co-cultured with KPC organoids for five days in normoxia. **(E)** Histogram of *IL6*-EGFP fluorescence intensity in PSCs. **(F)** Quantification of the relative MFI of *IL6*-EGFP in PSCs. N=3 biological replicates. Data represent mean+SD. P-values were calculated by two-way ANOVA. **(G)** Growth curve of tumors arising from subcutaneous co-injection of KPC cells alone or together with PSCs expressing control of *Vhl* sgRNA. N=9 mice. Data represent mean+/-SEM. P-values were calculated by two-way ANOVA.

## Discussion

Poor vascularization and the resulting generation of hypoxic areas are a feature of the microenvironment of virtually all solid tumors (38). In particular, hypoxia is has long been recognized as a characteristic of the PDAC TME and is associated with poor outcomes of PDAC patients which is at least in part due to its influence on the cancer cells (15,16,18,19). Whether hypoxia also affects the stromal cell state in the PDAC TME and their influence on tumor progression is less well understood. Here, we show that hypoxia shifts pancreatic fibroblasts towards acquisition of an inflammatory state that is tumor supporting. In addition, hypoxia potentiates the effects of cytokines secreted by PDAC cells that can promote the iCAF phenotype (10). This is consistent with our observations that iCAFs accumulate biochemical markers of hypoxia and that αSMA-negative CAFs are largely absent from hypoxic regions in murine PDAC. Furthermore, iCAFs display a hypoxic gene expression profile in human PDAC patients. These data indicate hypoxia as an environmental regulator of fibroblast heterogeneity in PDAC. Besides reduced oxygen tension, another consequence of inadequate vascularization and cancer cell metabolic activity in tumors is nutrient deprivation. In our *in vivo* and MEMIC experiments, we could not distinguish whether effects on αSMA or IL6 expression are mediated by hypoxia or nutrient scarcity or a combination thereof. However, our experiments in hypoxic culture conditions suggest that hypoxia alone can be sufficient to induce an inflammatory phenotype in PSCs.

The idea that hypoxia promotes an inflammatory response has been supported by several studies. In mice, short term exposure to hypoxia is sufficient to promote accumulation of inflammatory cells in several tissues and increases serum levels of various cytokines (39). In humans, three nights at high altitude increases levels of IL6 in the circulation (40). In addition to being observed in tumors, hypoxia is also a feature of wounds, and fibroblast heterogeneity has been observed in wound healing (41,42). Thus, our observations further support the idea that cancer cells can co-opt the normal stromal regenerative response to support tumor growth (43).

We found that the hypoxia-induced shift of PSCs towards an iCAF state is mediated by HIF-1α. Furthermore, hypoxia potentiates the ability of cytokines to promote acquisition of an iCAF phenotype in PSCs in a HIF-1α-dependent fashion. Elevated HIF-1α protein levels in hypoxic PSCs stimulated with cytokines likely results from increased *Hif1*_α_ transcription induced by NFkB signaling as a result of cytokine stimulation (34,44). Furthermore, cytokine signaling and HIF-1α cooperate to activate HIF-1α transcriptional activity by co-binding of STAT3 to promoter regions of HIF-1α target genes (45). Given that hypoxia can promote inflammatory cytokine production in cancer cells (46), a feed forward mechanism resulting in autocrine cytokine signaling is also conceivable. While our data indicate a major role of HIF-1α in the hypoxia-induced inflammatory response, we cannot exclude involvement of HIF-2α in this process. HIF-2α but not HIF-1α expression in αSMA^+^ CAFs has been shown to accelerate PDAC progression by establishing an immunosuppressive TME (47).

We provide evidence that *Vhl* deletion in PSCs induces an inflammatory response and promotes tumor growth in an allograft co-injection experiment. While the *in vitro* data suggests that it is the *Vhl*-deletion induced inflammatory cytokine expression that mediates this effect, we cannot exclude that other effects resulting from *Vhl* deletion in PSCs could support tumor growth. In addition, it is possible that other effects resulting from *Vhl* deletion besides the stabilization of HIF-1α are involved in this process.

Taken together, our work suggests that targeting tumor hypoxia could reduce accumulation of pro-tumorigenic iCAFs in PDAC and slow down tumor growth. Given the presence of hypoxia and CAF heterogeneity in most solid tumors (9,38), targeting hypoxic signaling in the tumor stroma might be a generalizable strategy to impair cancer progression.

## Methods

### Mouse experiments

All animal experiments described adhered to policies and practices approved by Memorial Sloan Kettering Cancer Center’s Institutional Animal Care and Use Committee (IACUC) and were conducted as per NIH guidelines for animal welfare (Protocol Number 11-03-007, Animal Welfare Assurance Number FW00004998). The maximal tumor size/burden permitted by the IACUC (Tumor burden may not exceed 10% of the weight of the mouse which is equivalent to a tumor volume of 2.5 cm^3^ for a 25 g mouse) was not exceeded. Mice were maintained under specific pathogen-free conditions and housed at 4-5 mice per cage at a 12-hour light/dark cycle at a relative humidity of 30% to 70% and room temperature of 22.2 ± 1.1°C, and were allowed access to food and water *ad libitum*. Mice were maintained in individually ventilated polysulfone cages with a stainless-steel wire bar lid and filter top on autoclaved aspen chip bedding. Mice were fed a closed-formula, natural-ingredient, γ-irradiated diet (5053 -PicoLab® Rodent Diet 20, Purina LabDiet) which was surface decontaminated using “flash” sterilization (100ºC for 1 minute). Mice were provided reverse-osmosis acidified (pH 2.5 to 2.8, with hydrochloric acid) water. Cage bottoms were changed weekly, whereas the wire bar lid, filter top and water bottle were changed biweekly.

### Orthotopic organoid injection

Orthotopic injections were performed as described (48). Organoids derived from pancreatic tumors of KPC (*Kras*^LSL-G12D/+^;*Trp53*^LSL-R172H/+^;Pdx1-Cre) in a C57BL/6 background were used. Syngeneic C57BL/6 mice were anesthetized with isoflurane and an incision was made in the left abdominal side. Organoids were dissociated from cultures with TrypLE (Thermo Fisher) and resuspended in 30 µL growth factor reduced Matrigel (Corning). Approximately 1×10^5^ cells were injected per recipient mouse into the tail region of the pancreas using a Hamilton Syringe. Successful injection was verified by the appearance of a fluid bubble without signs of intraperitoneal leakage. The abdominal wall was sutured with absorbable Vicryl sutures (Ethicon), and the skin was closed with wound clips (CellPoint Scientific Inc.). Mice were monitored for tumor development by ultrasound five weeks after injection and one/week afterwards using a Vevo 2100 System with a MS250 13-24MHz scan head (VisualSonics). When tumors were approximately 500 mm^3^ in size, 60 mg/kg body weight of pimonidazole (Hypoxyprobe) in 0.9% saline was injected i.p. one hour before euthanasia. Tumors were collected, and half of the tumor was allocated for 10% formalin fixation for histological analysis, and the other half was used to generate single cell suspensions for flow cytometry analysis.

### Immunofluorescence staining of mouse PDAC tumors

Automated multiplex IF was conducted using the Leica Bond BX staining system. Paraffin-embedded tissues were sectioned at 5 μm and baked at 58°C for 1 hr. Slides were loaded in Leica Bond and immunofluorescence staining was performed as follows. Samples were pretreated with EDTA-based epitope retrieval ER2 solution (AR9640, Leica) for 20 min at 95°C. The quadruplex-plex antibody staining and detection was conducted sequentially. The primary antibodies against PDPN (0.05 µg/ml, hamster, DSHB, 8.1.1), PIMO (0.12 µg/ml, mouse, Hydroxyprobe Inc. MAB1), SMA (0.1 µg/ml, rabbit, Abcam, ab5694) were used. For the rabbit antibody, Leica Bond Polymer anti-rabbit HRP was used, for the hamster antibody and the mouse antibody, rabbit anti-Hamster (Novex, A18891) and rabbit anti-mouse (Abcam, ab133469) secondary antibodies were used as linkers before the application of the Leica Bond Polymer anti-rabbit HRP. After that, Alexa Fluor tyramide signal amplification reagents (Life Technologies, B40953, B40958) or CF dye tyramide conjugates (Biotium, 92174, 96053) were used for detection. After each round of IF staining, Epitope retrieval was performed for denaturization of primary and secondary antibodies before another primary antibody was applied. After the run was finished, slides were washed in PBS and incubated in 5 μg/ml 4’,6-diamidino-2-phenylindole (DAPI) (Sigma Aldrich) in PBS for 5 min, rinsed in PBS, and mounted in Mowiol 4–88 (Calbiochem). Slides were kept overnight at -20°C before imaging.

### Imaging and analysis

Images from tissue sections of PDAC tumors were acquired with a Mirax Slide Scanner at 40x magnification. Images were analyzed in ImageJ. Pimonidazole^+^ regions were located in each tissue section. Within each region, the number of PDPN^+^ only pixels, SMA^+^ only pixels, and double positive pixels were quantified. Thresholds were set manually for each channel and kept consistent for each image. Two sections per tumor were analyzed. Live images from PSC monocultures were acquired with a Leica SP5 Inverted confocal microscope with cells placed in an environmental chamber.

### Cell culture

293T cells were obtained from ATCC (CRL-3216). PSCs were isolated from either wildtype C57BL/6 mice or *αSMA*-DsRed mice (29) by differential centrifugation as previously described (49) and immortalized by spontaneous outgrowth. Two lines of PSCs were used throughout the study. KPC (*Kras*^LSL-G12D/+^;*Trp53*^LSL-R172H/+^;Pdx1-Cre) mouse PDAC cells and organoids were described before (48). All cells were cultured at 37°C in 5% CO_2_ and 20% O_2_ and were maintained in DMEM supplemented with 10% FBS (Gemini), 100 U/ml penicillin and 100 µg/ml streptomycin (1% P/S). For hypoxia experiments, cells were cultured in a hypoxia chamber (Coy) set at 0.5% O_2_, 37°C and 5% CO_2_ for 48h. Cells were verified as mycoplasma-free by the MycoAlert Mycoplasma Detection Kit (Lonza). Cells were treated with 2 ng/mL murine IL1 (211-11A, Peprotech) and TNFα (315-01A, Peprotech) as indicated (“cytokines”).

### Organoid culture

Organoids were derived from pancreatic tumors of KPC (*Kras*^LSL-G12D/+^;*Trp53*^LSL-R172H/+^;Pdx1-Cre) mice in a C57BL/6 background and described before (48). Organoids were cultured 24-well plates in growth factor reduced (GFR) Matrigel (Corning) in advanced DMEM/F12 supplemented with the following: 1% P/S, 2 mM glutamine, 1X B27 supplement (12634-028, Invitrogen), 50 ng/ml murine EGF (PMG8043, Peprotech), 100 ng/ml murine Noggin (250-38; Peprotech), 100 ng/ml human FGF10 (100-26; Peprotech), 10 nM human Leu-Gastrin I (G9145, Sigma), 1.25 mM N-acetylcysteine (A9165; Sigma), 10 mM nicotinamide (N0636; Sigma), and R-spondin1 conditioned media (10% final). Organoids were passaged with every 3-4 days. For PSC co-culture, confluent wells of organoids were dissociated with 1x TrypLE (12604013, Thermo Fisher) and plated at a splitting ratio of 1:5 (approximately 1×10^4^ cells) together with 8×10^4^ *αSMA*-DsRed expressing PSCs in GFR Matrigel. Co-cultures were cultured with DMEM supplemented with 10% FBS (Gemini) and 1% P/S in 20% O_2_ and 5% CO_2_. For experiments in hypoxia, co-cultures were placed in a hypoxia chamber (Coy) set at 0.5% O_2_ for the last 48h of the experiment.

### MEMIC experiments

MEMICs were fabricated and used as described in detail previously (31). In brief, MEMICs were 3D printed in a 12-well format, and coverslips were glued at the bottom and the top to create inner and outer chambers. For each condition tested, one well was prepared without the coverslip on the top to create a control well without gradients. MEMICs were washed with water, UV-sterilized, washed twice with PBS and once with complete media before cell seeding. A 85 µL cell suspension containing 2×10^4^ PSCs was filled in the inner chamber. For the open wells, 1.5 mL of a 1×10^5^/mL cell suspension was added to the entire well. Cells were allowed to settle for 1h, and 1.5 mL media was added in the outer chamber in wells plated with cells in the inner chamber. The next day, cells were mock-treated or treated with cytokines. The gradient was allowed to form for 48h, and cells were either imaged live or fixed with 4% paraformaldehyde for 10 min, permeabilized with 0.1% Triton X-100, blocked with 2.5% bovine serum albumin in PBS, and stained for 1h with an anti-GFP antibody (A10262, Invitrogen). Wells were washed three times with PBS and incubated with an anti-chicken Alexa Fluor 488 coupled secondary antibody (A11039, Invitrogen) and Hoechst for nuclear staining for 30 min before being washed three times with PBS. Wells were imaged using BZ-X800 microscope from Keyence (20x magnification) and stitched using the BZ-X800 analysis software. Images were processed using custom MATLAB scripts. GFP/UnaG and DsRed fluorescence intensities were quantified and plotted according to their distance to the opening of the well. For *per* cell fluorescence quantification, images were segmented using nuclear staining and dilated to include adjacent cytoplasmic areas creating a mask for each cell. Then total fluorescence was integrated for each cell using these masks. Image analysis code is available upon request.

### Ectopic gene expression and CRISPR/Cas9 mediated gene deletion

Guide RNAs targeting murine *Hif1a and Vhl* were designed using GuideScan (http://www.guidescan.com/) and cloned into pLentiCRISPRv2 (Addgene 52961). The following guide sequences were used: TCGTTAGGCCCAGTGAGAAA (*Hif1a* sg1), CAAGATGTGAGCTCACATTG (*Hif1a* sg2), CCGATCTTACCACCGGGCAC (*Vhl* sg1), GGCTCGTACCTCGGTAGCTG (*Vhl* sg2). Rosa26 targeting guides (Ctrl sg) were described before (25). To create IL6 and hypoxia reporters, a Gibson assembly-based modular assembly platform (GMAP) was used (50). HRE-dUnaG from pLenti-HRE-dUnaG (Addgene 124372), and a PGK driven hygromycin selection cassette from MSCV Luciferase PGK-hygro (Addgene 18782) were amplified using primers containing overhangs with the homology sites for GMAP cloning and inserted into a lentiviral vector (LV 1-5; Addgene, 68411). IL6-EGFP from pmIL-6promoterEGFP (Addgene 112896), and a PGK driven blasticidin selection cassette from pMSCV-Blasticidin (Addgene 75085) were amplified for GMAP similarly and inserted into LV 1-5. Lentiviral particles were produced in 293T cells by using psPAX2 and pCMV-VSV-G packaging plasmids (Addgene 12260, 8454). Viral supernatant was collected after 48h, passed through a 0.45 µm nylon filter and used to transduce PSCs in the presence of 8 μg/mL polybrene (Sigma) overnight. Cells were subjected to puromycin (2 μg/mL, Sigma), hygromycin (250 μg/mL) or blasticidin (10 μg/mL, Invivogen) antibiotic selection the following day. Polyclonal cell populations were used for the experiments.

### Western blot

Lysates were generated by incubating cells in RIPA buffer (Millipore). 20-30 μg of cleared lysate were analyzed by SDS-PAGE as previously described (25). The following primary antibodies were used: Vinculin (1:5000 dilution, Sigma, V9131), β-Actin (1:5000; Sigma, A5441), HIF-1α (1:1000, 10006421, Cayman), VHL (1:200, sc-5575, Santa Cruz), LDHA (1:1000, 2012S, Cell Signaling) and GFP (1:1000, 11814460001, Sigma). The following secondary antibodies were used: anti-rabbit HRP (1:5000, NA934V, GE) and anti-mouse HRP (1:5000, NA931, GE).

### Flow cytometry

For analysis of PSC monocultures, cells were trypsinized, washed, stained with DAPI and analyzed on an LSRFortessa II (BD). Live cells (DAPI-) were analyzed for EGFP fluorescence. For organoid/PSC co-cultures, Matrigel was digested with Dispase (Corning), and cells and organoids were dissociated mechanically by pipetting up and down at least 30 times. PSCs were analyzed by gating for DAPI- and DsRed+ cells followed by analysis of EGFP fluorescence intensity. For analysis of CAFs from PDAC tumors arising from orthotopic injection of KPC organoids, tumors were minced and resuspended in 5 mL DMEM with 800 µg/mL Dispase (Sigma), 500 µg/mL Collagenase P (Sigma), 100 ug/mL Liberase TL (Roche), 100 µg/mL DNaseI (Sigma), 100 µg/mL Hyaluronidase (Sigma). Samples were then transferred to C-tubes and processed using program 37C_m_TDK1_2 on a gentleMACS Octo dissociator with heaters (Miltenyi Biotec). Dissociated tissue was passaged through a 40 µm cell strainer and centrifuged at 1500 rpm x 5 minutes. Red blood cells were lysed with ACK lysis buffer (A1049201, Thermo Fisher) for 1 minute, and tubes were filled up with PBS. Samples were centrifuged and resuspended in FACS buffer (PBS supplemented with 2% FBS) and stained with Ghost Dye Violet 510 (1:1000, Tonbo Biosciences) on ice for 10 min for discrimination of viable and non-viable cells. Samples were blocked with anti-CD16/32 (FC block, 1:100, Biolegend) for 15 minutes on ice and then incubated with the following antibodies (all from Biolegend) in Brilliant stain buffer (Thermo Fisher) for 30 minutes on ice: CD326-FITC (G8.8, 1:50), CD45-BV711 (30-F11, 1:200), CD31-PE/Cy7 (390, 1:200), PDPN-APC/Cy7 (8.1.1, 1:100), Ly6C-BV421 (HK1.4, 1:200). Samples were washed in FACS buffer and fixed and permeabilized with the Foxp3 / Transcription Factor Staining Buffer Set (00-5523-00, Thermo Fisher) according to the manufacturer’s instructions. Samples were stained with anti-pimonidazole antibody (4.3.11.3, 1:50, Hypoxyprobe) in permeabilization buffer at 4ºC over night. Samples were incubated with anti-mouse Alexa Fluor 647 (1:400, Thermo Fisher) in permeabilization buffer for 15 min at room temperature. Samples were resuspended in FACS buffer and analyzed on an LSRFortessa II by gating for Ghost Dye-, CD45-, CD31, CD326-, PDPN+ cells comparing Ly6C-with Ly6C+ cells. Compensation was performed with UltraComp eBeads (01-2222-42, Thermo Fisher). Data were analyzed with FlowJo software (BD).

### Quantification of gene expression

Total RNA was isolated from fibroblasts with Trizol (Life Technologies) according to the manufacturer’s instructions, and 1 µg RNA was used for cDNA synthesis using iScript (Bio-Rad). Quantitative real-time PCR (qPCR) analysis was performed in technical triplicates using 1:20 diluted cDNAs and 0.1 µM forward and reverse primers together with Power SYBR Green (Life Technologies) in a QuantStudio 7 Flex (Applied Biosystems). Gene expression was quantified in Microsoft Excel 365 as relative expression ratio using primer efficiencies calculated by a relative standard curve. The geometric mean of the endogenous control genes *Actb* and *Rplp0* was used as reference sample. Primer pairs are as follows: TACCACCATGTACCCAGGCA (*Actb* FW), CTCAGGAGGAGCAATGATCTTGAT (*Actb* RV), AGATTCGGGATATGCTGTTGGC (*Rplp0* FW), TCGGGTCCTAGACCAGTGTTC (*Rplp0* RV), CCATCATGCGTCTGGACTT (*αSMA* FW), GGCAGTAGTCACGAAGGAATAG (*αSMA* RV), CTTCCATCCAGTTGCCTTCT (*IL6* FW), CTCCGACTTGTGAAGTGGTATAG (*IL6* RV), GTGTCAACCACTGTGCTAGT (*Cxcl1* FW), CACACATGTCCTCACCCTAATAC (*Cxcl1* RV), CATTGTCAAGTACAGTCCACACT (*Ldha* FW), TTCCAATTACTCGGTTTTTGGGA (*Ldha* RV).

### RNA sequencing

Total RNA was isolated with Trizol as above, and libraries were prepared from polyA-selected mRNA using the TruSeq RNAsample preparation kit v2 (Illumina) according to the manufacturer’s instructions. Libraries were sequenced using an Illumina HiSeq 4000 generating 150 bp paired-end reads. An average of 58 million reads per sample was retrieved. Adaptor sequences were removed from fastq files with Trimmomatic v.0.36, and trimmed reads were mapped to the mus musculus GRCm38 reference genome using the STAR aligner v.2.5.2b. Aligned features were counted with featureCounts from the Subread package v.1.5.2 and differential expression was determined using DESeq2 v3.10 from Bioconductor in R v4.1.0.

### Gene set enrichment analysis (GSEA)

GSEA was performed using a pre-ranked gene list based on the log2 fold change comparing two Ctrl sg samples cultured in Normoxia against two Ctrl sg samples cultured in Hypoxia for 48h, or comparing two Ctrl sg samples cultured in Hypoxia against two *Hif1a* sg7 samples cultured in Hypoxia. GSEA 4.3.0 (Broad Institute) was used with 1000 permutations and mouse gene symbols remapped to human orthologs v7.5 (MSigDB). Enrichment of the iCAF signature (4) or Hallmark signatures (MSigDB) was analyzed.

### Statistics

A student’s *t*-test was applied to compare one variable between two groups. One-way ANOVA was applied to compare one variable between three or more groups. Two-way ANOVA was applied to compare two independent variables between two groups. Correction for multiple comparisons was done using the Holm-Sidak method. Statistical analysis was done in GraphPad Prism 9. Most graphs show the mean + SD with individual datapoints, unless indicated otherwise in the figure legends.

## Acknowledgments

We thank the members of the Thompson laboratory for helpful discussions. We are thankful to Tullia Lindsten for help with planning of and protocol preparation for mouse experiments, and to Natalya N. Pavlova for help with GMAP. We also thank Elisa De Stanchina and Inna Kudos from the MSKCC Antitumor Assessment Core for help with orthotopic organoid injections into the pancreas, and Wenfei Kang and Eric Rosiek of the MSKCC Molecular Cytology Core for help with immunofluorescence staining, microscopy and image analysis. S.S. receives support from the NCI (1K99CA259224) and the Alan and Sandra Gerry Metastasis and Tumor Ecosystems Center at MSKCC. S.S. was also supported by the Human Frontier Science Program (LT000854/2018). K.M.T. was supported by the Jane Coffin Childs Memorial Fund for Medical Research and a Shulamit Katzman Endowed Postdoctoral Research Fellowship. This work was supported by MSKCC’s David Rubenstein Center for Pancreatic Research Pilot Project (to S.W.L.) and NIH grant P01CA013106 (to S.W.L.). S.W.L. is an Investigator of the Howard Hughes Medical Institute and the Geoffrey Beene Chair for Cancer Biology at MSKCC. C.C.F. receives support from the National Cancer Institute at the NIH (DP2 CA250005), the American Cancer Society (RSG-21-179-01-TBE) and the Pew Charitable Trust (00034121). C.B.T. was supported by the NCI (R01CA201318). This work used core facilities at MSKCC that were supported by the cancer center support grant (P30CA008748).

## Author Contributions

S.S. conceived the project, performed most experiments, analyzed data, interpreted results, and wrote and edited the manuscript. M.R. performed, analyzed and interpreted MEMIC experiments. K.M.T. assisted with experiments using KPC organoids, ultrasound monitoring and collection of PDAC tumors. F.V.C. provided technical assistance. S.W.L. and C.C.F. provided support for PDAC and MEMIC experiments, respectively. C.B.T. interpreted results, and wrote and edited the manuscript. All authors participated in discussing and finalizing the manuscript.

---

**Supplementary Figure 1.**
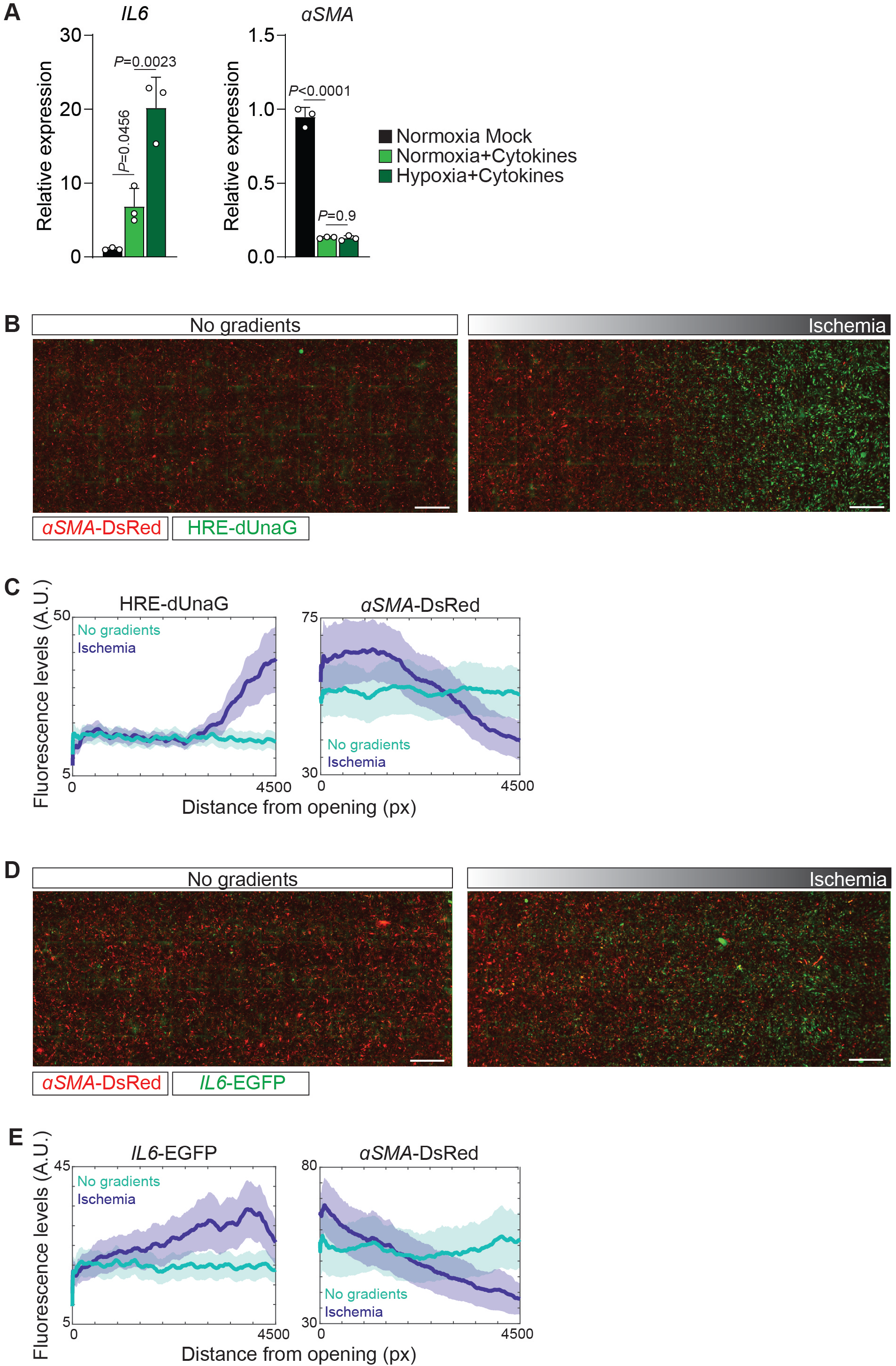
**(A)** qPCR for the indicated transcripts in PSCs cultured in Normoxia and mock-treated or treated with cytokines (IL1/TNFα), or cultured in Hypoxia and treated with cytokines for 48h. N=3 biological replicates. Data represent mean+SD. P-values were calculated by one-way ANOVA. **(B, C)** MEMIC experiments with hypoxia reporter. **(B)** Representative images of PSCs expressing *αSMA*-DsRed and HRE-dUnaG, treated with cytokines and cultured in the MEMIC without a cover (no gradients, left) or with a cover (ischemia, right) for 48h. Scale bar = 500 µm. **(C)** Quantification of HRE-dUnaG (left) and *αSMA*-DsRed (right) fluorescence intensity with increasing distance from the oxygen-rich opening. A.U. = arbitrary units, px = pixel. **(D, E)** MEMIC experiments with IL6 reporter. **(D)** Representative images of PSCs expressing *αSMA*-DsRed and *IL6*-EGFP, treated with cytokines and cultured in the MEMIC without a cover (no gradients, left) or with a cover (ischemia, right) for 48h. Scale bar = 500 µm. **(E)** Quantification of *IL6*-EGFP (left) and *αSMA*-DsRed (right) fluorescence intensity with increasing distance from the oxygen-rich opening.

**Supplementary Figure 2.**
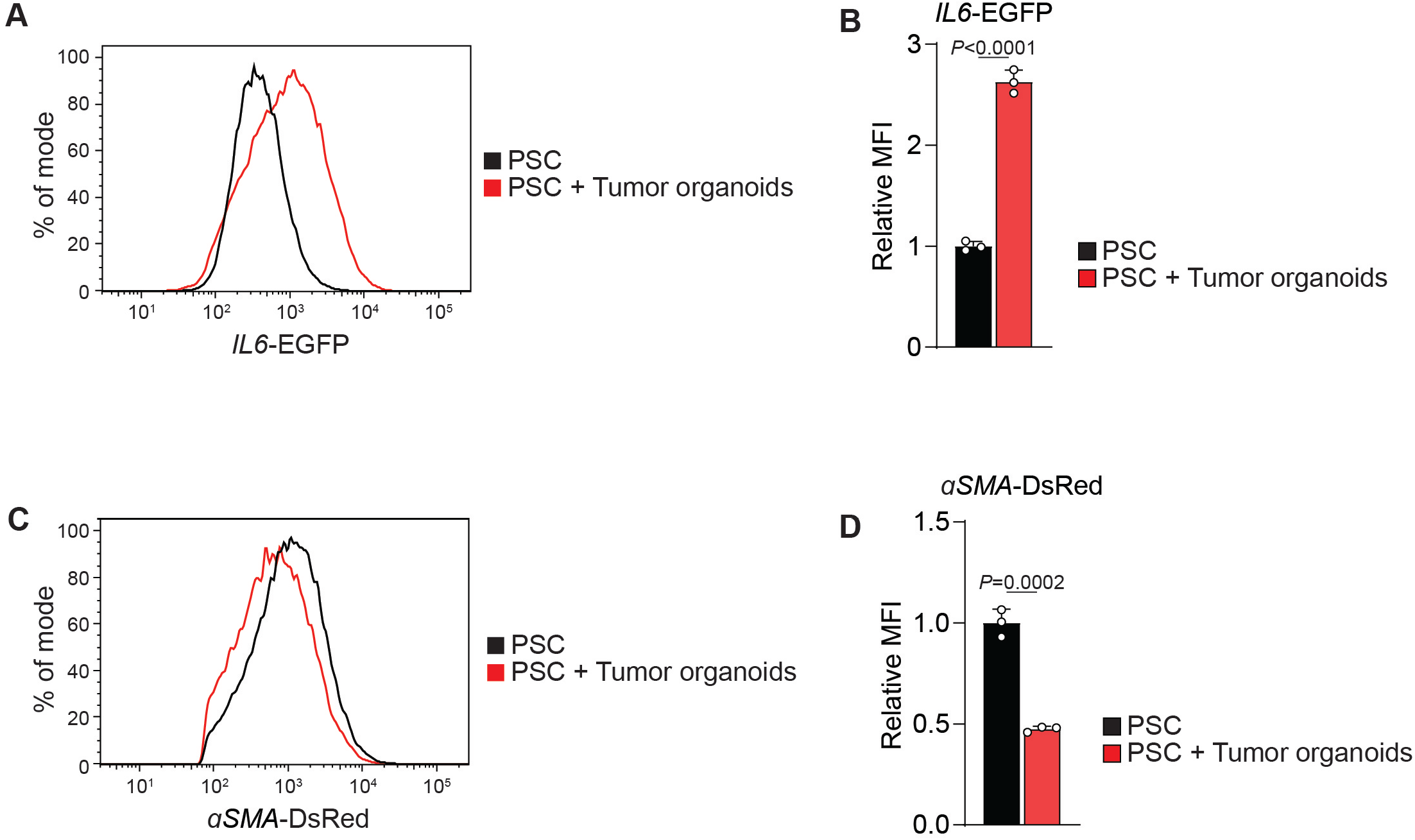
**(A-D)** PSC/Tumor organoid co-culture experiment. PSCs expressing *IL6*-EGFP and αSMA-DsRed were cultured alone or together with KPC organoids for five days. **(A)** Histogram of *IL6*-EGFP fluorescence intensity in PSCs. **(B)** Quantification of the relative MFI of *IL6-*EGFP in PSCs. **(C)** Histogram of *αSMA*-DsRed fluorescence intensity in PSCs. **(D)** Quantification of the relative MFI of *αSMA*-DsRed in PSCs. N=3 biological replicates. Data represent mean+SD. P-values were calculated by Student’s t-test.

**Supplementary Figure 3.**
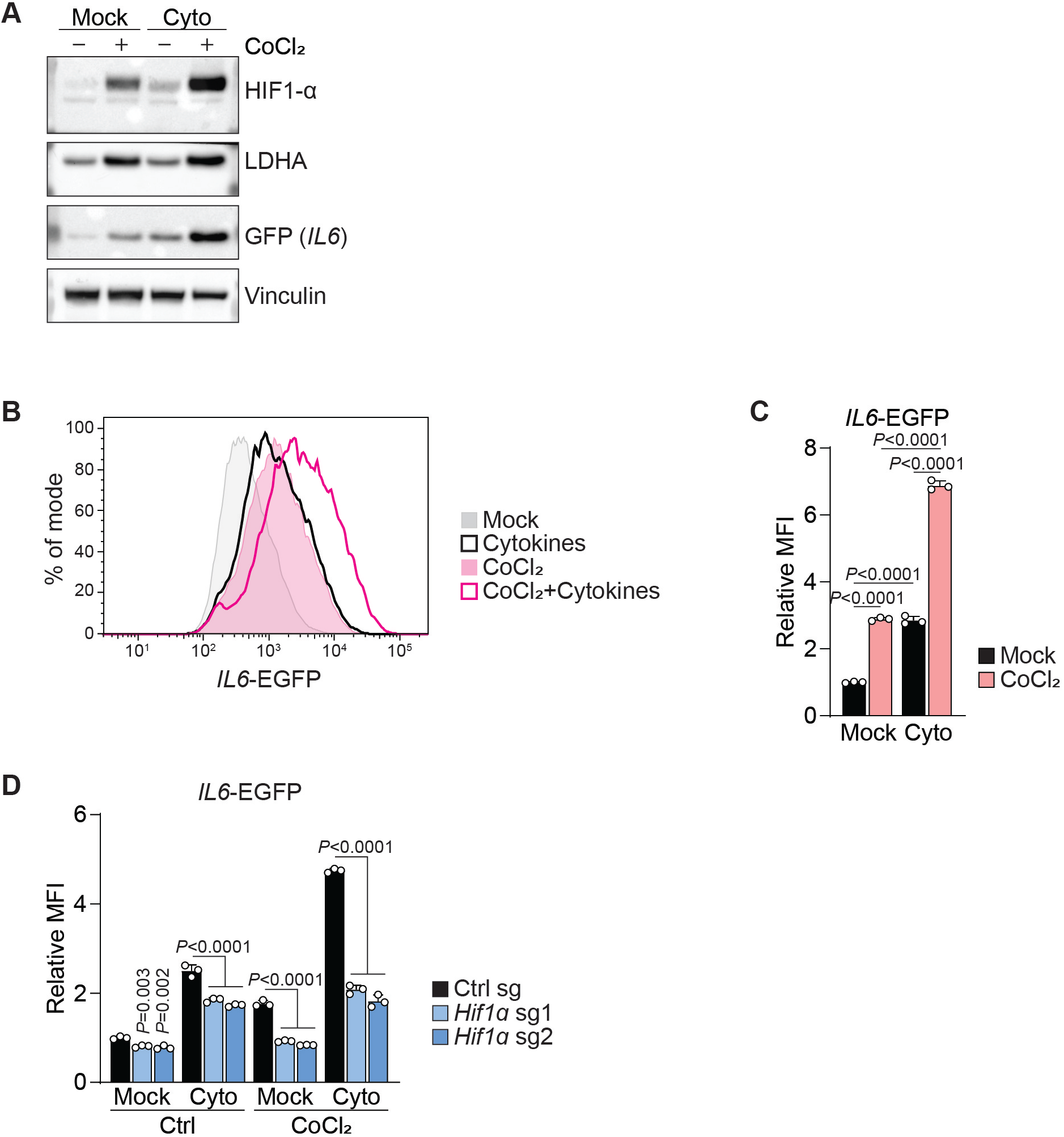
**(A)** Western blot of PSCs expressing *IL6*-EGFP that were mock treated, treated with 100 µM CoCl_2_, cytokines or a combination thereof for 48h. **(B, C)** Fluorescence intensity of *IL6*-EGFP expressing PSCs that were mock treated or treated with 100 µM CoCl_2_, cytokines or a combination thereof for 48h. **(B)** Histogram of *IL6*-EGFP fluorescence intensity. **(C)** Quantification of relative MFI of *IL6*-EGFP. N=3 biological replicates. Data represent mean+SD. P-values were calculated by two-way ANOVA. **(D)** Quantification of relative MFI in PSCs expressing *IL6*-EGFP and control or *Hif1a* sgRNA that were mock treated or treated with 100 µM CoCl_2_, cytokines or a combination thereof for 48h. N=3 biological replicates. Data represent mean+SD. P-values were calculated by two-way ANOVA.

